# Improvements to Casanovo, a deep learning *de novo* peptide sequencer

**DOI:** 10.1101/2025.07.25.666826

**Authors:** Gwenneth Straub, Varun Ananth, William E. Fondrie, Chris Hsu, Daniela Klaproth-Andrade, Michael Riffle, Justin Sanders, Bo Wen, Lingwen Xu, Melih Yilmaz, Michael J. MacCoss, Sewoong Oh, Wout Bittremieux, William Stafford Noble

## Abstract

Casanovo is a state-of-the-art deep learning model for *de novo* peptide sequencing from mass spectrometry proteomics data. Here we report on a series of enhancements to Casanovo, aimed at improving the interpretability of the scores assigned to predicted peptides, generalizing the software for use in database search, speeding up training and prediction runtimes, and providing workflows and visualization tools to facilitate adoption of Casanovo and interpretation of its results. Our goal is to make Casanovo accurate and easy to use for applications such as metaproteomics, antibody sequencing, immunopeptidomics, and discovery of novel peptide sequences in standard proteomics analyses. Casanovo is available as open source at https://github.com/Noble-Lab/casanovo.

## Introduction

Most tandem mass spectrometry (MS/MS)-based proteomics data are initially interpreted using a database search procedure that aims to infer peptide sequences from the observed mass spectra. For spectra generated via data-dependent acquisition, this mapping is ideally one-to-one: each spectrum corresponds to a single generating peptide. The database search procedure combs through a given list of potential generating peptides and finds the one peptide that best explains each observed spectrum. The database search paradigm works well when a high quality list of potential generating peptides can be easily assembled. This is the case, for example, when the sample comes from a single organism whose complete genome has been sequenced. However, in some settings, such as metaproteomics or antibody sequencing, the list of potential peptides is not easily available. In these cases, *de novo* sequencing is required.

The first algorithms for *de novo* sequencing of peptides from tandem mass spectra were proposed forty years ago [1], and dozens of new algorithms have been proposed since then (reviewed in [2, 3]). The field advanced considerably in 2015 when machine learning was first applied to this problem [4], and then advanced again in 2017 with the first deep learning model for *de novo* sequencing, DeepNovo [5].

Casanovo [6], introduced in 2022, was the first deep learning model to employ a transformer architecture for *de novo* sequencing. Transformers were initially developed for natural language processing [7] and have subsequently been adapted to many different application domains, including modeling DNA and protein sequences [8, 9]. Most prior models for *de novo* sequencing required discretization of the *m/z* axis, thereby yielding long, sparse vectors or requiring a substantial loss in *m/z* precision. In contrast, the transformer can handle variable-length inputs, so each mass spectrum can be natively represented as a list of (*m/z*, intensity) tuples. Indeed, the transformer architecture has been so successful at *de novo* sequencing that most subsequently developed models have adopted this framework [10].

In practice, to be useful to the proteomics community, a *de novo* sequencing method needs to perform well on real data and also be available as easy-to-use software with broad functionality. In this work, we describe several substantial enhancements to Casanovo, which are implemented in version 5.0 of the software. These improvements include a modified peptide scoring procedure that yields much better calibrated confidence estimates, a substantial speedup in the software for training and inference, a new command that allows Casanovo’s learned score function to be used for database search, and containerization and visualization tools that make Casanovo easier to use and its outputs easier to interpret.

## Methods

### Casanovo overview

The Casanovo model employs an encoder–decoder framework, where both the encoder and decoder are implemented using transformers (Figure 1A). The input to the model is an MS/MS spectrum, which is represented as a set of (*m/z*, intensity) tuples. This spectrum is processed by a multi-layer transformer (the encoder), which uses its built-in attention mechanism to identify important peak-to-peak relationships. The output of the transformer is a latent embedding of the spectrum. This embedding, along with the precursor mass and charge, is then provided as input to the decoder, which uses a transformer coupled with a beam search algorithm to iteratively predict each amino acid in the sequence, conditioning each prediction on the preceding predictions.

**Figure 1.**
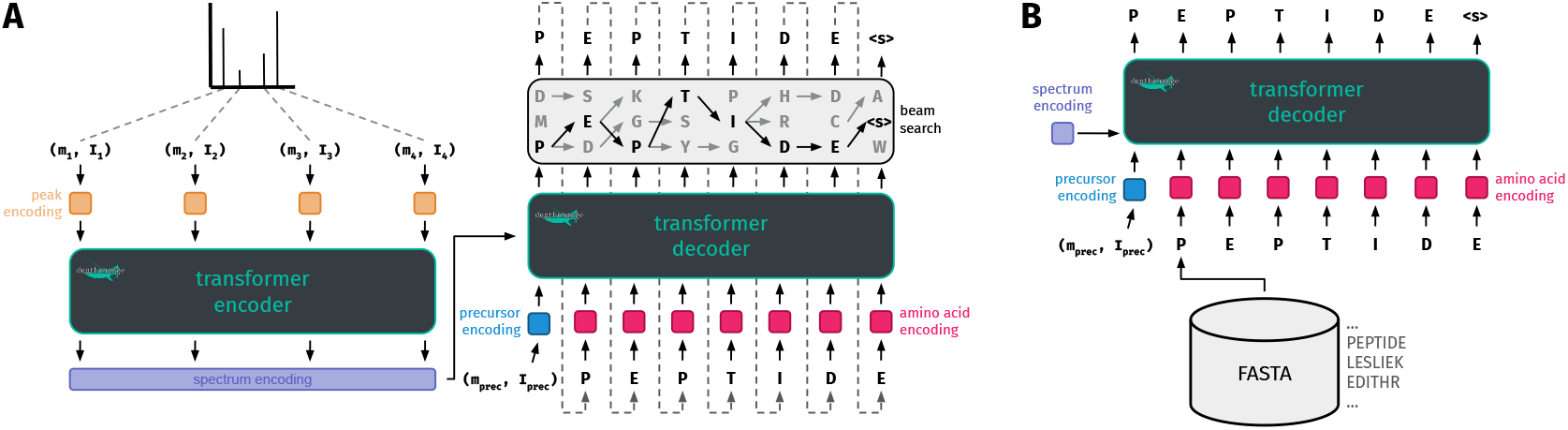
Casanovo neural network architecture. (A) Casanovo consists of an encoder–decoder neural network architecture. The encoder receives an MS/MS spectrum as input and converts it into a latent embedding. The decoder takes as input the spectrum embedding and precursor information, and autoregressively predicts the peptide sequence, in combination with beam search decoding. (B) During database searching, instead of autoregressive predicting, the decoder receives candidate peptide sequences and produces a PSM score using the teacher forcing method.

The Casanovo software operates in two modes. In training mode, the input is a collection of annotated spectra in structured Mascot Generic Format (MGF) files, in which each spectrum is associated with a peptide sequence and corresponding charge state. During training, the model weights are iteratively adjusted to encourage the model to predict the correct peptide for each observed spectrum in the training set. The output of this procedure is a weights file containing the learned parameters of the transformer model, which can be used to generate predictions. Subsequently, when Casanovo is used in sequencing mode, the weights file is provided as input, along with unknown spectra in MGF, mzML [11], or mzXML [12] files. In this mode, Casanovo’s weights do not change, and an inference algorithm uses the model to identify the best-scoring peptide sequence for each observed MS/MS spectrum. Hence, the output from the sequencing operation is a list of spectrum annotations, where each spectrum has a predicted peptide and an associated confidence score, output in the mzTab format [13]. In practice, Casanovo reports peptide-level confidence scores as well as scores for each predicted amino acid. Optionally, if annotated spectra are provided as input to the sequencing mode, it can be used to calculate various performance measures, such as peptide and amino acid level precision–coverage metrics.

### Recent enhancements to Casanovo

Under the hood, Casanovo is powered by a general Python library for modeling mass spectrometry data— both mass spectral data and peptide or small molecule analytes—called DepthCharge (https://github.com/wfondrie/depthcharge). DepthCharge provides the infrastructure that Casanovo builds upon to read mass spectrometry data files, through libraries such as Pyteomics [14], as well as encode the mass spectra and analytes for subsequent modeling. Since our earlier versions of Casanovo, DepthCharge has been upgraded for improved efficiency, including improvements in its internal data storage. Specifically, previous versions of DepthCharge used HDF5 [15] as internal format, whereas this has now been upgraded to the Lance file format (https://lancedb.github.io/lance), which is an Apache Arrow-based format that is built for fast random access of batched queries. Furthermore, DepthCharge now supports streaming mass spectra from their original mzML, MGF, or mzXML files, rather than first being cached. Casanovo benefits from these two changes during both training and sequencing for improved runtime performance. Numerous other small changes also benefit Casanovo: all mass calculations have been vectorized and can be performed on the same device as where the model is running, such as graphics processing units (GPUs); the intensity values of mass peaks are now optionally encoded using a sinusoidal waveform rather than merely a linear projection; mass spectrum preprocessing has been unified with spectrum utils [16]; and there is now first-class support for the ProForma 2.0 specification of peptides [17]. These upgrades make Casanovo consistent with Proteomics Standards Initiative standards [18] and, as we demonstrate below, substantially more efficient.

In addition to the DepthCharge upgrade, we have updated Casanovo’s beam search implementation to make better use of GPU parallelization and carry out more efficient PyTorch tensor operations. Casanovo’s beam search algorithm is employed during inference to decode peptide sequences. Decoding starts at the first residue position by selecting the top *k* scoring residues, where *k* is the user-specified number of beams (default = 1). At each subsequent step, the algorithm expands each beam by selecting its top *k* scoring next residues, resulting in *k*^2^ candidate beams. From these candidates, the top *k* scoring candidates are selected to proceed to the next decoding step. Decoding ends once all *k* beams have been terminated, which occurs when either a stop token is predicted or the predicted peptide sequence’s mass is greater than the precursor mass plus a configurable tolerance. We previously implemented much of this decoding algorithm directly in Python. However, standard Python operations cannot benefit from hardware acceleration and also suffer from the overhead of the Python interpreter. Therefore, we reimplemented the decoding algorithm to make use of PyTorch vectorized tensor operations, which can be executed largely outside of the Python interpreter and benefit from hardware acceleration.

In parallel with the efficiency upgrades, we have revamped the Casanovo command line interface to improve how input and output files are handled:

- Previously, Casanovo had an `eval` command to compute performance statistics relative to a given gold standard annotation. This functionality is now implemented as an option (`--evaluate`) on the `sequence` command.
- During training, Casanovo now stores the best model, i.e., the model with the lowest loss on the validation set, in an output file named `best.ckpt`. Previously, a set of top-performing models were stored, and the optimal one could only be identified by examining the associated log file.
- Output files for all commands can be placed in a specific directory using `--output dir` and using a particular filename prefix using `--output root`.
- A Boolean `--overwrite` option was added, which defaults to “False,” to prevent Casanovo from inadvertently overwriting files from a previous run.
- Training and validation loss values are now stored in a tab-delimited text file, for use in downstream analysis. Previously, these values were only available by parsing the Casanovo log file.
- The mzTab output files are automatically validated using the validator at https://apps.lifs-tools.org/mztabvalidator/ to ensure syntactic correctness [19].

In addition, we have modified how Casanovo’s peptide-level and amino acid-level scores are calculated. During inference, Casanovo autoregressively generates a peptide one amino acid at a time from the C-terminus to the N-terminus, and each amino acid is assigned a corresponding score by the model. To compute the peptide-level score, previous versions of Casanovo computed the arithmetic mean of these amino acid-level scores [20]. In the new version of Casanovo, we instead report the product of the amino acid-level scores. Furthermore, while the previous versions computed the score for each individual amino acid as the mean of the model-reported amino acid score and the peptide score, in the new version of Casanovo the model-reported amino acid scores are used directly, without averaging with the total peptide score.

### Database search

We have introduced a new database search mode in Casanovo, using the `db_search` command. In this mode, Casanovo uses knowledge gained from *de novo* sequencing in order to score PSMs in a database search paradigm [21]. Each mass spectrum is encoded into a latent representation by Casanovo’s encoder, and in a similar manner to the “teacher forcing” strategy used during training [22], Casanovo’s decoder takes the encoded mass spectrum and a candidate peptide as input and scores each amino acid, conditioned on the encoded mass spectrum and the previous amino acids in the candidate sequence (Figure 1B). The final peptide score is calculated by taking the product of these amino acid scores.

During database search, a target–decoy strategy is typically used to calculate false discovery rates (FDRs) [23]. Because Casanovo currently cannot directly generate decoy peptides necessary for this procedure, instead we used the `generate-peptides` command from Crux (version 4.3) [24] with default parameters to generate shuffled decoy peptides from a reference proteome. We then combined these decoy peptides with target peptides derived from the original reference proteome as input to Casanovo. Finally, we used Crema (with default parameters; version 0.0.10) [25] to calculate the FDR at the peptide level based on the Casanovo search results matching both target and decoy peptides.

### Data

To train and evaluate the newest Casanovo version, we used a previously described set of train, validation, and test splits [26]. These splits were originally drawn from peptide-spectrum matches (PSMs) from the MassIVE knowledge base (MassIVE-KB) [27], including 2 million tryptic PSMs from MassIVE-KB v2018-06-15 and *∗*1.1 million multi-enzyme PSMs from MassIVE-KB v2.0.15 (annotated as “proteomics experiments digested with various different enzymes”). Critically, the MassIVE-KB train, validation, and test sets are disjoint at the peptide level. More details about the creation of these splits can be found in **(author?)** [26]. To ensure an even distribution of digestion enzymes across the training set, we combined and shuffled the tryptic and multi-enzyme training data.

To demonstrate and benchmark Casanovo’s database search functionality, we used three MS/MS runs measured from human, mouse, and yeast samples, respectively. The samples were prepared using a protein aggregate method as described previously [28] and acquired on an Orbitrap Fusion Lumos mass spectrometer paired to an EvoSep One liquid chromatography system using an 88-minute extended gradient method. For all samples, the data-dependent acquisition method contained one cycle of MS1 (60,000 Orbitrap resolving power, 118 milliseconds max injection time, and 100% AGC target) and MS2 (15,000 Orbitrap resolving power, 1.6 *m/z* isolation window, 27% HCD collision energy, 22 milliseconds max injection time, and 100% AGC target). The MS1 peaks were filtered by intensity threshold of greater than 2E4, charge state of 2–5, and dynamic exclusion (with an exclusion duration of 15 seconds and repeat count of 1). Raw MS/MS data were converted to mzML files using msconvert with peak picking enabled in ProteoWizard (version 3.0.24031) [29]. Reference databases for each experiment were downloaded from UniProt [30] (proteome identifiers UP000005640, UP000000589, and UP001165141) on 18 April 2025.

## Results

### Casanovo produces well-calibrated confidence scores

An ideal *de novo* sequencing method not only produces accurate predictions but also provides an accompanying score that discriminates between correct and incorrect predictions. This score is even more useful if it has well-defined semantics. For example, if the score ranges from 0 to 1, then we might prefer that a score of 0.9 implies that the prediction has a 90% probability of being correct. Such a score is considered well *calibrated* [31]. Modifying the peptide-level score to use the product of the amino acid scores, rather than the arithmetic mean, yielded a substantial improvement in the calibration of the peptide scores. Intuitively, this approach makes sense if we interpret the amino acid scores assigned by the model as conditional likelihoods.

To test the calibration empirically, we applied Casanovo to a test set of 106,933 PSMs from the MassIVE-KB repository. We then sorted Casanovo’s predictions by score and computed, in a sliding window of size 501, the percentage of correct predictions. Plotting this empirical accuracy versus the Casanovo score at the center of the window yields a calibration curve. We observe that switching from the arithmetic mean to the product dramatically improves Casanovo’s overall calibration (Figure 2A). In this curve, the median difference between the empirical accuracy and the Casanovo score is 0.028, and the maximum deviation is 0.106. The largest discrepancies occur in the high-score range, where Casanovo’s score is somewhat conservative, in the sense that, e.g., a score of 0.90 corresponds to an empirical accuracy of 97%. We also evaluated the impact of this change in scoring scheme on Casanovo’s performance, observing that the two scoring schemes perform nearly identically, with an average precision of 80.86% for the arithmetic mean and 80.87% for the product (Figure 2B). Overall, reporting the peptide-level score as the product of the raw amino acid confidence scores provides superior calibration while having a negligible impact on the average precision.

**Figure 2.**
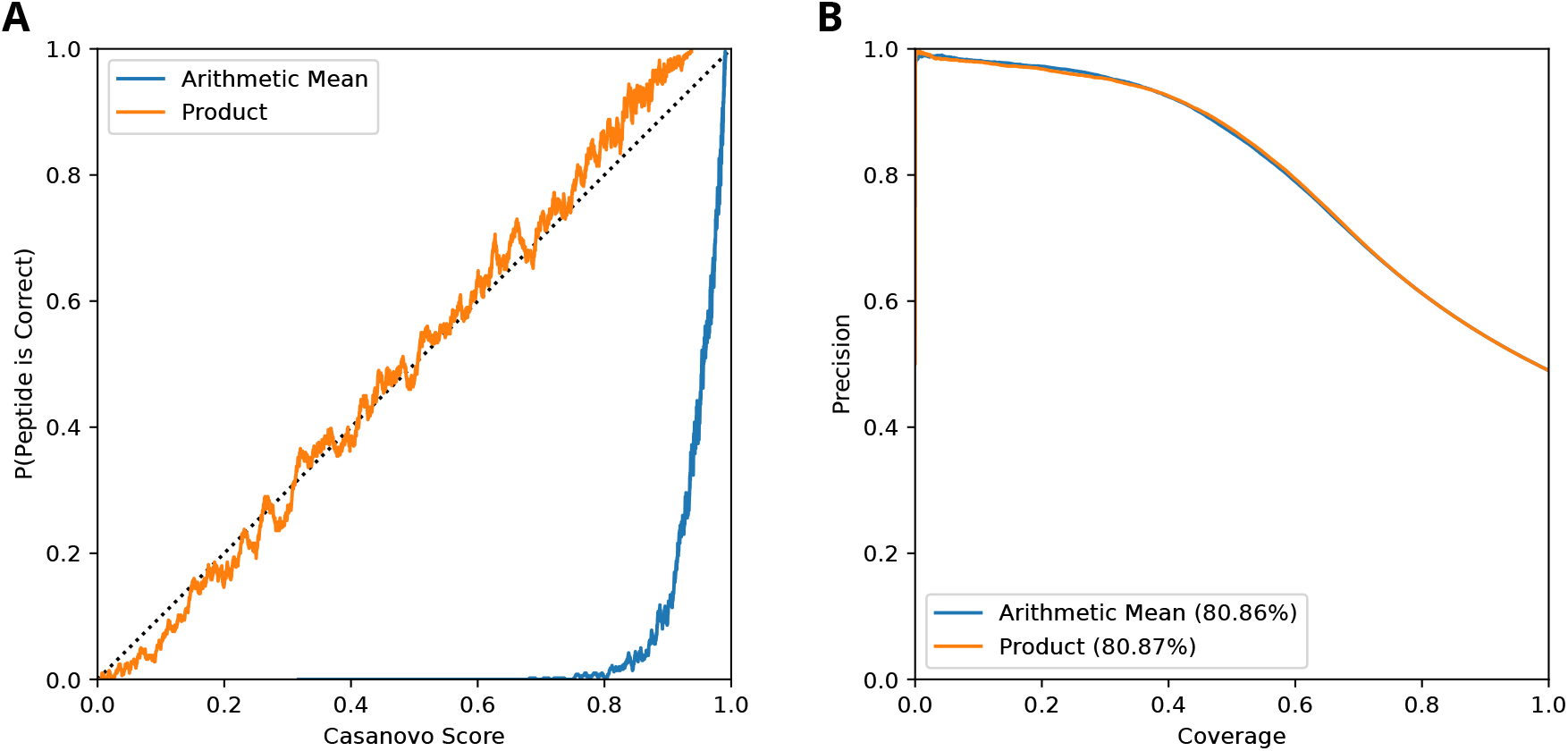
Casanovo peptide scores are well calibrated. (A) The proportion of correct predictions as a function of the Casanovo score for predictions on the MassIVE-KB test set. The percentages of correct predictions are computed in a sliding window of size 501 with a step size of 1. (B) Peptide-level precision as a function of coverage for two variants of the Casanovo scoring scheme (arithmetic mean and product of amino acid scores) on the MassIVE-KB test set. Values in the legend represent average precision.

### Casanovo can be used for database search

In previous work, we have demonstrated that a trained Casanovo model can be deployed not only for *de novo* sequencing but also as a score function for database search [21], and that Casanovo’s score function yields improved statistical power to detect peptides compared to hand-designed score functions such as XCorr [32], hyperscore [33], or the Andromeda score [34]. Previously, users interested in employing Casanovo in this way had to check out a separate branch on git, install the package from source locally, and use an outdated version of the Casanovo model. Casanovo v5.0 now provides this database search functionality in a new mode, `db_search`. Hence, a command of the form `casanovo db_search myspectra.mzml mydb.fasta` will instruct Casanovo to search the given spectra against the specified database, reporting the results in an mzTab file. This new functionality significantly improves the method’s accessibility.

In order to demonstrate and benchmark Casanovo’s new database search functionality, we compare Casanovo’s database search scores against XCorr, the score function proposed in the very first database search algorithm SEQUEST [32]. We use the XCorr implementation in the Tide search engine (version 4.3) [35], and we compare Casanovo and Tide results using three mass spectrometry runs, derived from human, mouse, and yeast samples. For both Tide and Casanovo we used the default set of parameters except for the minimum peptide length, which was set to 8. Because Casanovo was not trained to distinguish between leucine and isoleucine, we swapped all isoleucines with leucines within all three of the reference protein databases. As a performance measure, we compare the number of peptides detected across a range of peptide-level FDR thresholds.

This experiment confirms that Casanovo yields markedly improved statistical power to detect peptides, relative to Tide. In particular, at a 1% peptide-level FDR threshold, we observe an increase in the number of detected peptides of 61.3%, 57.1%, and 82.3% for the human, mouse, and yeast runs, respectively (Figure 3).

**Figure 3.**
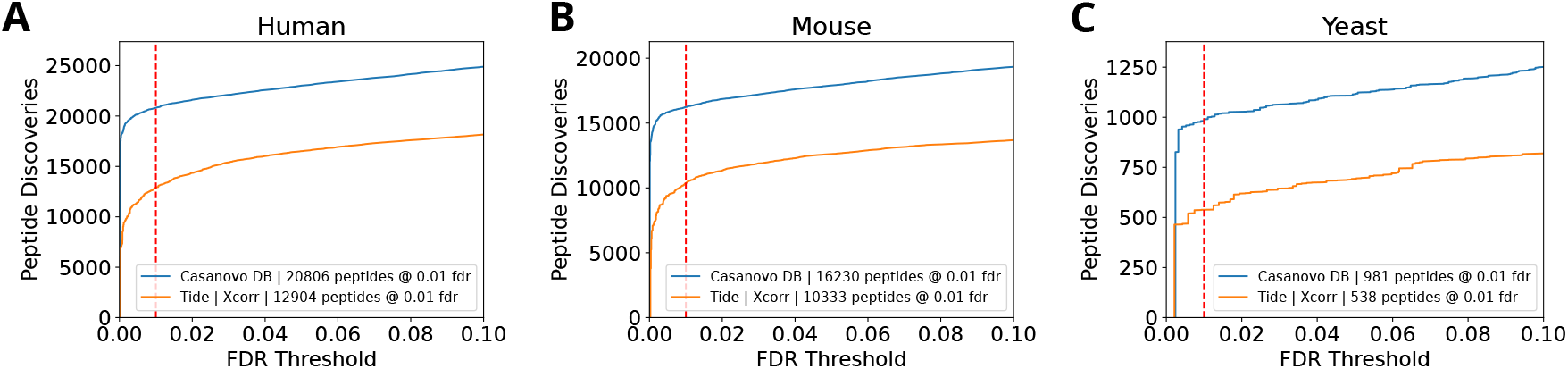
Comparing Casanovo’s learned score function with XCorr. Each panel plots, for the human (A), mouse (B), and yeast (C) experiments, the number of distinct peptides detected as a function of the peptide-level FDR threshold. The two series in each panel correspond to database search carried out using Casanovo and Tide, with the number of accepted peptides at 1% FDR listed in the legends.

### Casanovo is faster than it used to be

In order to calculate the relative speedup between the previous and new versions of Casanovo, for both versions we recorded the wall clock execution time for one epoch of training over Casanovo’s MassIVE-KB training set and five inference runs with a varying number of beams over Casanovo’s evaluation set. Both experiments were conducted in isolation on a machine with an Intel Xeon Gold 6426Y CPU, 1024 GB of RAM, and using an NVIDIA L40 48GB GPU. To ensure that both runs had equal access to system resources, such as the file system and memory caches, during both experiments the training and evaluation run was the only user (non-system) process running on the machine.

The results of this experiment suggest that Casanovo v5.0 achieves an 11.0-fold training mode speedup and a 3.5-fold inference mode speedup relative to v4.3 (Figure 4A). In practice, this means that, with our hardware setup, one epoch of training on a dataset of 2,896,373 spectra required 74,818 s using Casanovo v4.3 and now only requires 6,774 s with v5.0. Similarly, the inference speed increased from 76.3 spectra/s to 235 spectra/s.

**Figure 4.**
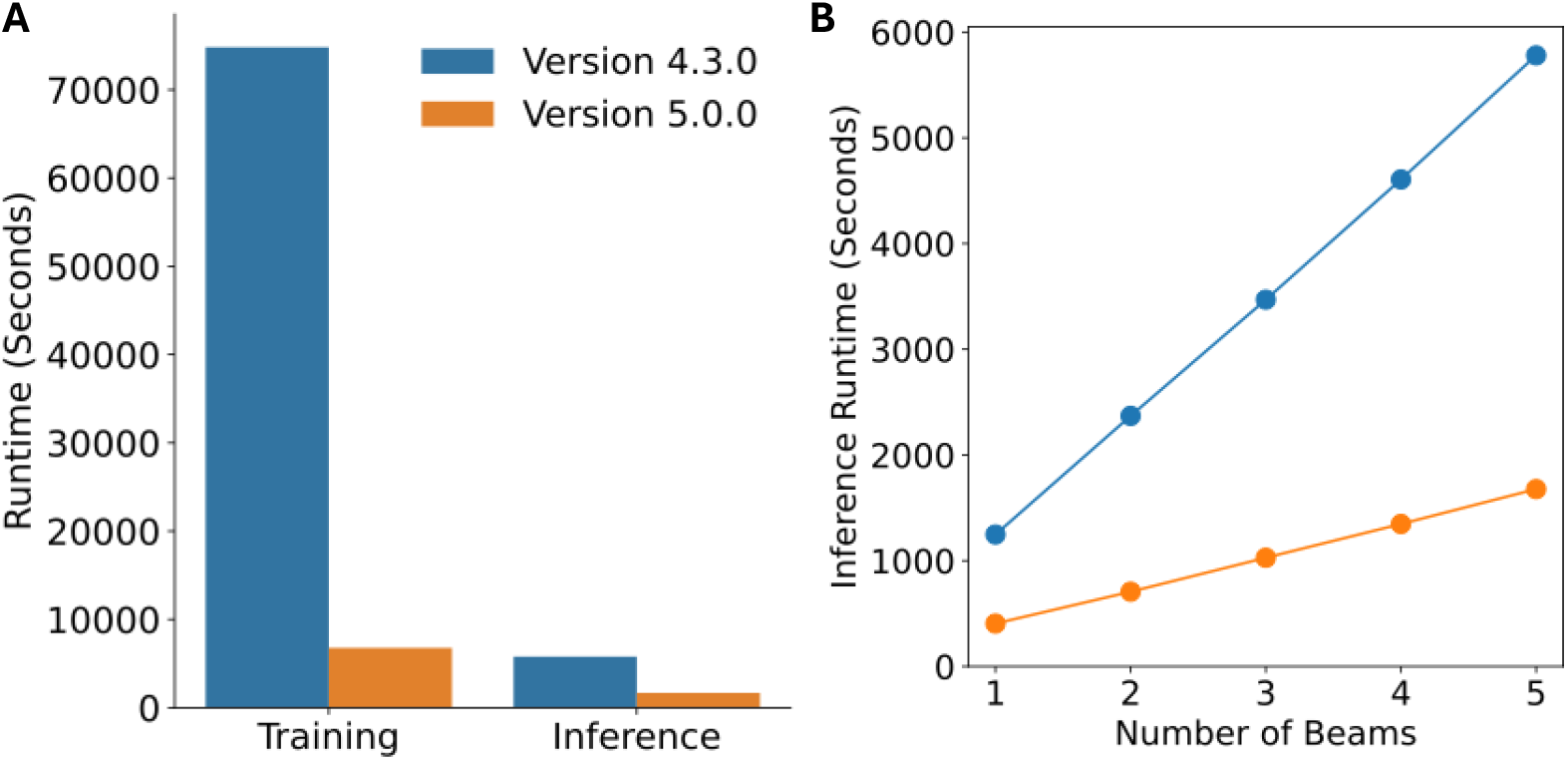
Casanovo runtime comparison. (A) Timing comparison of Casanovo v4.3 and v5.0 for training and inference using default parameters. Each bar corresponds to one pass of training over Casanovo’s training data or inference over Casanovo’s evaluation data. (B) Inference runtime over Casanovo’s evaluation data as a function of the number of beams.

Most of the speedup to Casanovo’s inference mode is driven by the upgrades we made to the beam search implementation. To further quantify this speedup, we benchmarked inference for Casanovo v4.3 and v5.0 varying the number of beams from one through five. This experiment showed that the new beam search implementation achieves a 3.07-fold speedup when using one beam and a 3.46-fold speedup when using five beams (Figure 4B). In practice, Casanovo v5.0 inference on this dataset can be done on our hardware at a rate of 234 spectra/s using one beam and 56.7 spectra/s using five beams.

### Tools for easily running Casanovo and visualizing its results

#### Visualization using PDV

PDV [39] (https://github.com/wenbostar/PDV) allows users to visualize PSMs generated from a variety of proteomics search engines. The tool runs on a local computer through a graphical user interface and is capable of exporting publication-quality annotated spectra. PDV has been extended to support visualizing Casanovo results, thereby allowing users to evaluate how well a spectrum aligns with the peptide predicted by Casanovo. Users can also create “mirror spectrum” visualizations to easily compare Casanovo’s alignment with one obtained using an alternative peptide (Figure 5A). Finally, to further assess the confidence of the Casanovo prediction, users can compare with the match between the predicted peptide and its predicted spectrum using deep learning models such as Prosit [40] and AlphaPeptDeep [41] (Figure 5B). These predictions are generated by PDV using a publicly available Koina server [42]. A step-by-step tutorial on how to use PDV to visualize Casanovo results is available at https://github.com/wenbostar/PDV/wiki/Visualize-Casanovo-result.

**Figure 5.**
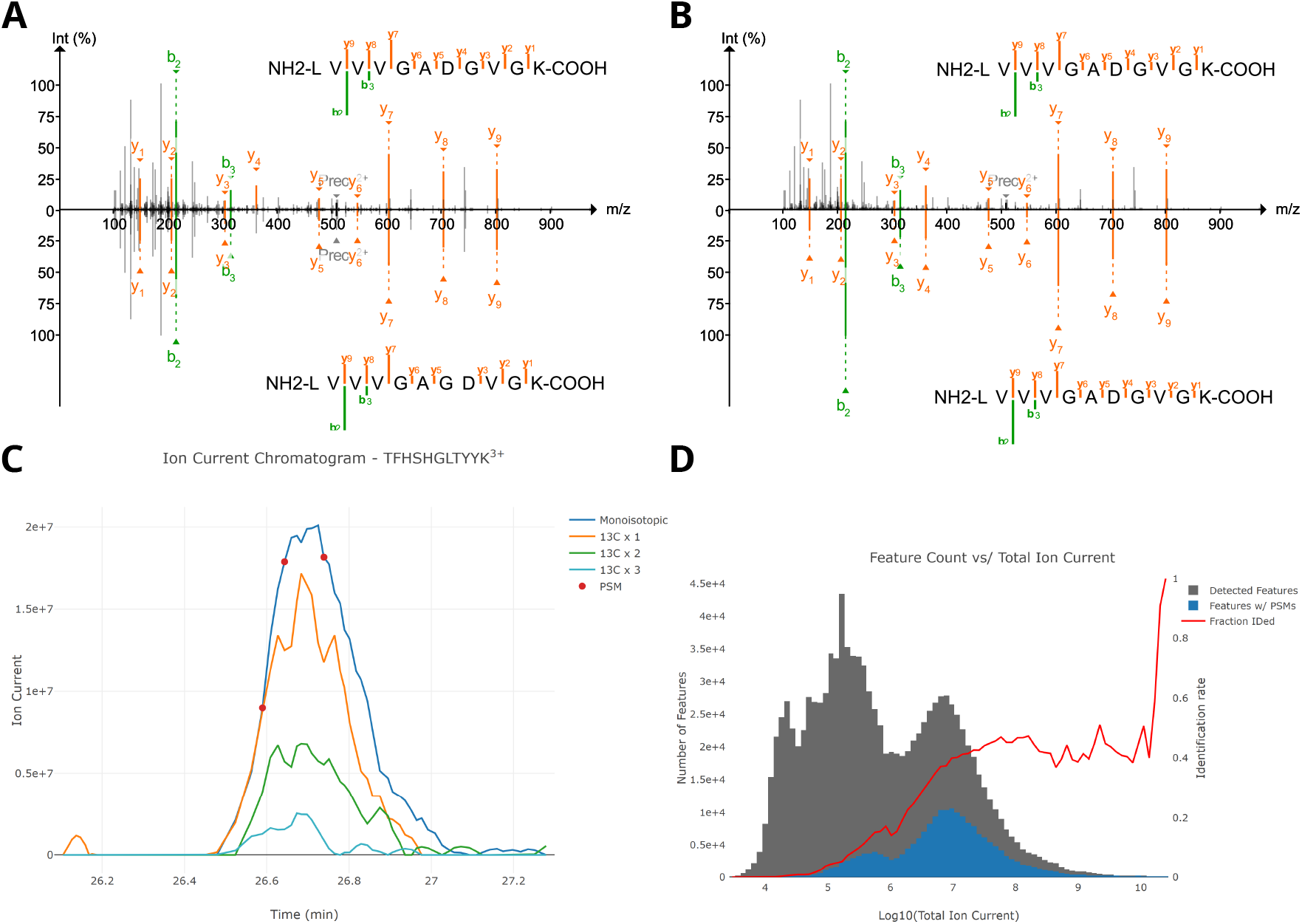
Visualizing Casanovo results. (A) PDV visualization of two different annotations for a given spectrum. The top panel shows a detection for KRAS G12D mutant peptide using Casanovo. This PSM was reported in a previous study using a peptide-centric search method, and the detection was supported by genomics data [36]. The bottom panel shows the annotation for the same spectrum using a different mutant peptide from the same gene (KRAS G13D). The comparison shows that the G12D peptide is a better match for the spectrum than the G13D peptide. (B) PDV visualization comparing an observed spectrum to a spectrum with fragment ion intensities predicted by AlphaPeptDeep. The top PSM is the same as (A), but the bottom PSM shows the spectrum predicted by AlphaPeptDeep. (C) An example extracted ion chromatogram of MS1 signal generated by Limelight for a Casanovo search result. The colored lines represent the intensity of the monoisotopic and +1, +2, and +3 13C isotopic precursor ions over time. The red circles indicate where Casanovo identified a PSM for this peptide ion. (D) An example of a quality control chart generated by Limelight. The gray histogram shows the number of peptide features predicted by Hardklor [37] and Bullseye [38] with the given intensities. The blue histogram shows the number of these predicted features identified by Casanovo. The red line shows the fraction of features at those intensities that were identified by Casanovo. The red line indicates that more intense features are more likely to be identified by Casanovo, as expected.

#### Visualization, annotation, and sharing of results with Limelight

Limelight [43] is a generalized web application for viewing, analyzing, and sharing data-dependent acquisition proteomics data. We have extended Limelight to enable analysis of Casanovo results. To upload results to Limelight, users first convert Casanovo results to Limelight XML using the converter available at https://github.com/yeastrc/limelight-import-casanovo. The resulting Limelight XML can be manually uploaded to a Limelight project using the data upload interface within Limelight. More information about uploading data and using Limelight may be found here: https://limelight-ms.readthedocs.io/.

Limelight’s design is agnostic with regard to data analysis workflow, so once Casanovo data are in Limelight, all of its features can be applied to Casanovo’s *de novo* search results. This functionality includes advanced sharing tools and access to the full data stack, including annotated MS and MS/MS spectra, on-demand extracted ion chromatograms (Figure 5C), peptide- and PSM-based views of search results, and many quality control (QC) visualizations (Figure 5D). Users can compare search hits from Casanovo to other search programs, including comparing results scan-by-scan, comparing peptide lists, and visually comparing QC metrics.

#### Containerization: Docker and Nextflow

We have containerized Casanovo using Docker, which creates a distributable image that includes Casanovo and the entire environment necessary for Casanovo to run. We have uploaded the Casanovo Docker image to Docker Hub, and it is available at `mriffle/casanovo:5.0.0`. This image allows users to run Casanovo without needing to explictly install Casanovo or any of its prequisites. Once Docker is installed, users can run the latest version of Casanovo via the command `docker run --rm -it --shm-size=1g mriffle/casanovo:5.0.0 casanovo.` For full documentation of the Docker image, see https://github.com/mriffle/casanovo-docker.

We leveraged the Docker image above to develop a workflow using Nextflow [44] that greatly simplifies running an entire *de novo* data analysis using Casanovo. The workflow allows users to run all steps of a peptide search workflow, including RAW file conversion to mzML, analysis with Casanovo, and upload to Limelight. Users do not need to install any of the components of the workflow (e.g., msconvert or Casanovo)—all components are run using Docker images and Nextflow orchestrates the running of each step. The steps can be run on a user’s computer, a computer cluster, or on the cloud with Amazon Web Services. To run the workflow, users need only install Nextflow, though to run analyses locally, users also need to install Docker. We have documented the installation and running of the workflow here: https://nf-ms-dda-casanovo.readthedocs.io/.

## Discussion

Overall, we hope that the improvements described here to Casanovo enhance its utility to the scientific community. In this update, we have addressed several practical limitations by improving usability, efficiency, and interoperability, making Casanovo easier to adopt and integrate into proteomics workflows. Version 5.0 reflects a significant step forward in software maturity: it provides better-calibrated confidence scores, faster runtime performance, a streamlined interface for both *de novo* and database search, and integration with visualization and sharing platforms to support broader adoption.

The calibration results reported above are useful, in the sense that they provide an intuitive way to set a threshold on Casanovo’s scores. On the other hand, there is still work to be done before we can accurately control the FDR in a *de novo* setting. Formally, controlling the FDR requires proving that the proportion of false positives is bounded in expectation, i.e., on average in a statistical sense. In principle, this proportion can be computed by summing over the posterior error probability of each prediction in a ranked list, up to the specified threshold. However, by definition, our calibration results are only computed using labeled spectra to which peptides could be confidently assigned by database searching. In contrast, Casanovo is applied to all spectra in a given run at inference time, not just those that are identifiable. Thus, the FDR needs to be controlled in a more realistic collection of both high quality and lower quality spectra. Additionally, because Casanovo was trained using teacher forcing, from a theoretical standpoint we would expect its learned amino acid scores to be well calibrated only when conditioned on a correct peptide prefix. Thus, as is a common challenge in autoregressive models, accumulation of errors may hurt calibration and hinder any efforts to use scores directly for FDR control. Finally, users typically are interested primarily in discovering *de novo* predictions that are not in the reference proteome. Therefore, methods to control the FDR specifically among this subset of discoveries, as opposed to all predictions, are required.

We also caution that, currently, Casanovo’s db search mode should still be considered experimental, in the sense that we have not yet succeeded in making the code fast enough to be practically useful in most settings. For example, analyzing the yeast dataset required 16 hours and 4 minutes of wall clock time, which is 264 times longer than the same analysis using Tide. We hypothesize that a model distillation procedure, in which we optimize a much smaller model to approximate Casanovo’s scores, could dramatically reduce this running time.

Taken together, these enhancements improve the reliability, efficiency, and interpretability of Casanovo’s predictions. We anticipate that these developments will help support more confident use of *de novo* sequencing in practical proteomics workflows and provide a solid foundation for future extensions of the tool and its applications.

## Data availability

The train, validation, and test splits utilized for training and benchmarking Casanovo are available on Zenodo at https://zenodo.org/records/12587317. In particular, we used the splits found under the directory `multi_enzyme_simple.` The human, mouse, and yeast MS/MS data is available at https://panoramaweb.org/Casanovo_v5.url with ProteomeXChange identifier PXD066485. The reference human, mouse, and yeast proteomes used in the database search experiments can be downloaded from UniProt at https://www.uniprot.org/proteomes/UP000005640, https://www.uniprot.org/proteomes/UP000000589, and https://www.uniprot.org/proteomes/UP001165141, respectively.

## Software availability

The open-source Casanovo software is available with an Apache 2.0 license on GitHub at https://github.com/Noble-Lab/casanovo/ and is mirrored on Zenodo at 10.5281/zenodo.11205038.

## Acknowledgments

This work was funded by National Science Foundation award 2245300 to WSN, by the National Science Foundation Graduate Research Fellowship Program (Grant No. DGE-2140004, B.W.), and by the University of Washington’s Proteomics Resource (UWPR95794). DK-A acknowledges funding from the Deutsche Forschungsgemeinschaft (DFG, German Research Foundation) via the IT Infrastructure for Computational Molecular Medicine (#461264291) and through the project CLINSPECT-M (16LW0243K). WB acknowledges support by the University of Antwerp Research Fund, the University of Antwerp Industrial Research Fund, and the Research Foundation – Flanders (FWO G087625N).

## Author contributions

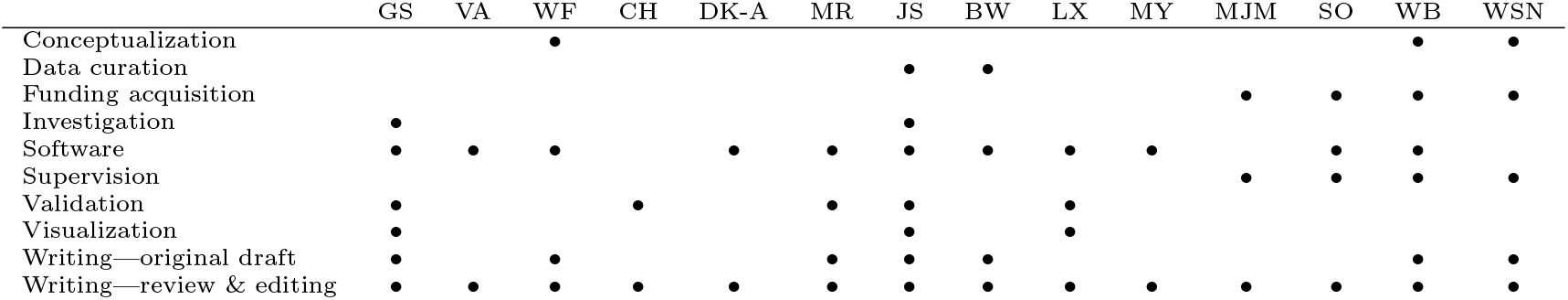

## References

[1] T. Sakurai, T. Matsuo, H. Matsuda, and I. Katakuse. Paas 3: A computer program to determine probable sequence of peptides from mass spectrometric data. Biomedical Mass Spectrometry, 11(8):396– 399, 1984.

[2] Rui Vitorino, Sofia Guedes, Fabio Trindade, Inês Correia, Gabriela Moura, Paulo Carvalho, Manuel AS Santos, and Francisco Amado. De novo sequencing of proteins by mass spectrometry. Expert Review of Proteomics, 17(7-8):595–607, 2020.

[3] Cheuk Chi A Ng, Yin Zhou, and Zhong-Ping Yao. Algorithms for de-novo sequencing of peptides by tandem mass spectrometry: A review. Analytica Chimica Acta, page 341330, 2023.

[4] B. Ma. Novor: Real-time peptide de novo sequencing software. Journal of the American Society for Mass Spectrometry, 26:1885–1894, 2015.

[5] N. H. Tran, X. Zhang, L. Xin, B. Shan, and M. Li. De novo peptide sequencing by deep learning. Proceedings of the National Academy of Sciences of the United States of America, 31:8247–8252, 2017.

[6] M. Yilmaz, W. E. Fondrie, W. Bittremieux, S. Oh, and W. S. Noble. De novo mass spectrometry peptide sequencing with a transformer model. In Proceedings of the International Conference on Machine Learning, pages 25514–25522, 2022.

[7] A. Vaswani, N. Shazeer, N. Parmar, J. Uszkoreit, L. Jones, A. N. Gomez, L. Kaiser, and I. Polosukhin. Attention is all you need. Advances in Neural Information Processing Systems, 30, 2017.

[8] A. Rives, J. Meier, T. Sercu, S. Goyal, Z. Lin, J. Liu, D. Guo, M. Ott, C. L. Zitnick, J. Ma, and R Fergus. Biological structure and function emerge from scaling unsupervised learning to 250 million protein sequences. Proceedings of the National Academy of Sciences of the United States of America, 118(15):e2016239118, 2021.

[9] Žiga Avsec, Vikram Agarwal, Daniel Visentin, Joseph R Ledsam, Agnieszka Grabska-Barwinska, Kyle R Taylor, Yannis Assael, John Jumper, Pushmeet Kohli, and David R Kelley. Effective gene expression prediction from sequence by integrating long-range interactions. Nature Methods, 18(10):1196–1203, 2021.

[10] W. Bittremieux, V. Ananth, W. Fondrie, C. Melendez, M. Pominova, J. Sanders, B. Wen, M. Yilmaz, and W. S. Noble. Deep learning methods for de novo peptide sequencing. Mass Spectrometry Reviews, 2024. In press.

[11] L. Martens, M. Chambers, M. Sturm, D. Kessner, F. Levander, J. Shofstahl, W. H. Tang, A. Rompp, S. Neumann, A. D. Pizarro, L. Montecchi-Palazzi, N. Tasman, M. Coleman, F. Reisinger, P. Souda, H. Hermjakob, P. A. Binz, and E. W. Deutsch. mzML–a community standard for mass spectrometry data. Mol Cell Proteomics, 10(1):R110.000133, Ja. 2011.

[12] Patrick GA Pedrioli, Jimmy K Eng, Robert Hubley, Mathijs Vogelzang, Eric W Deutsch, Brian Raught, Brian Pratt, Erik Nilsson, Ruth H Angeletti, Rolf Apweiler, et al. A common open representation of mass spectrometry data and its application to proteomics research. Nature biotechnology, 22(11):1459–1466, 2004.

[13] J. Griss, A. R. Jones, T. Sachsenberg, M. Walzer, L. Gatto, J. Hartler, G. G. Thallinger, R. M. Salek, C. Steinbeck, N. Neuhauser, et al. The mzTab data exchange format: communicating mass-spectrometry-based proteomics and metabolomics experimental results to a wider audience. Molecular and Cellular Proteomics, 13(10):2765–2775, 2014.

[14] Anton A Goloborodko, Lev I Levitsky, Mark V Ivanov, and Mikhail V Gorshkov. Pyteomics—a Python framework for exploratory data analysis and rapid software prototyping in proteomics. Journal of The American Society for Mass Spectrometry, 24(2):301–304, 2013.

[15] Quincey Koziol, Russ Rew, Mark Howison, et al. Hdf5. In High Performance Parallel I/O, pages 227–244. Chapman and Hall/CRC, 2014.

[16] Wout Bittremieux, Lev Levitsky, Matteo Pilz, Timo Sachsenberg, Florian Huber, Mingxun Wang, and Pieter C. Dorrestein. Unified and standardized mass spectrometry data processing in python using spectrum utils. Journal of Proteome Research, 22(2):625–631, Januar. 2023.

[17] Richard D LeDuc, Veit Schwammle, Michael R Shortreed, Anthony J Cesnik, Stefan K Solntsev, Jared B Shaw, Maria J Martin, Juan A Vizcaino, Emanuele Alpi, Paul Danis, et al. ProForma: a standard proteoform notation. Journal of Proteome Research, 17(3):1321–1325, 2018.

[18] Eric W Deutsch, Juan Antonio Vizcaíno, Andrew R Jones, Pierre-Alain Binz, Henry Lam, Joshua Klein, Wout Bittremieux, Yasset Perez-Riverol, David L Tabb, Mathias Walzer, et al. Proteomics standards initiative at twenty years: current activities and future work. Journal of Proteome Research, 22(2):287–301, 2023.

[19] Nils Hoffmann, Joel Rein, Timo Sachsenberg, Jürgen Hartler, Kenneth Haug, Gerhard Mayer, Oliver Alka, Saravanan Dayalan, Jake TM Pearce, Philippe Rocca-Serra, et al. mzTab-M: a data standard for sharing quantitative results in mass spectrometry metabolomics. Analytical Chemistry, 91(5):3302–3310, 2019.

[20] M. Yilmaz, W. E. Fondrie, W. Bittremieux, R. Nelson, V. Ananth, S. Oh, and W. S. Noble. Sequence-to-sequence translation from mass spectra to peptides with a transformer model. Nature Communications, 15(1):6427, 2024.

[21] V. Ananth, J. Sanders, M. Yilmaz, B. Wen, S. Oh, and W. S. Noble. A learned score function improves the power of mass spectrometry database search. Bioinformatics (ISMB), 2024. In press.

[22] R. J. Williams and D. Zipser. A learning algorithm for continually running fully recurrent neural networks. Neural Computation, 1(2):270–280, 1989.

[23] J. E. Elias and S. P. Gygi. Target-decoy search strategy for increased confidence in large-scale protein identifications by mass spectrometry. Nature Methods, 4(3):207–214, 2007.

[24] C. Y. Park, A. A. Klammer, L. Käll, M. P. MacCoss, and W. S. Noble. Rapid and accurate peptide identification from tandem mass spectra. Journal of Proteome Research, 7(7):3022–3027, 2008.

[25] A. Lin, D. See, W. E. Fondrie, U. Keich, and W. S. Noble. Target-decoy false discovery rate estimation using Crema. Proteomics, page 2300084, 2023.

[26] Carlo Melendez, Justin Sanders, Melih Yilmaz, Wout Bittremieux, Will Fondrie, Sewoong Oh, and William Stafford Noble. Accounting for digestion enzyme bias in casanovo. Journal of Proteome Research, 23(10):4761–4769, 2024.

[27] M. Wang, J. Wang, J. Carver, B. S. Pullman, S. W. Cha, and N. Bandeira. Assembling the community-scale discoverable human proteome. Cell Systems, 7:412–421.e5, 2018.

[28] Bo Wen, Chris Hsu, Wen-Feng Zeng, Michael Riffle, Alexis Chang, Miranda Mudge, Brook L Nunn, Matthew D Berg, Judit Villen, Michael J MacCoss, et al. Carafe enables high quality in silico spectral library generation for data-independent acquisition proteomics. bioRxiv, pages 2024–10, 2024.

[29] M. C. Chambers, B. Maclean, R. Burke, D. Amodei, D. L. Ruderman, S. Neumann, L. Gatto, B. Fischer, B. Pratt, J. Egertson, K. Hoff, D. Kessner, N. Tasman, N. Shulman, B. Frewen, T. A. Baker, M. Y. Brusniak, C. Paulse, D. Creasy, L. Flashner, K. Kani, C. Moulding, S. L. Seymour, L. M. Nuwaysir, B. Lefebvre, F. Kuhlmann, J. Roark, P. Rainer, S. Detlev, T. Hemenway, A. Huhmer, J. Langridge, B. Connolly, T. Chadick, K. Holly, J. Eckels, E. W. Deutsch, R. L. Moritz, J. E. Katz, D. B. Agus, M. J. MacCoss, D. L. Tabb, and P. Mallick. A cross-platform toolkit for mass spectrometry and proteomics. Nature Biotechnology, 30(10):918–920, 2012.

[30] UniProt Consortium. UniProt: a hub for protein information. Nucleic Acids Research, page gku989, 2014.

[31] U. Keich and W. S. Noble. On the importance of well calibrated scores for identifying shotgun proteomics spectra. Journal of Proteome Research, 14(2):1147–1160, 2015.

[32] J. K. Eng, A. L. McCormack, and J. R. Yates, III. An approach to correlate tandem mass spectral data of peptides with amino acid sequences in a protein database. Journal of the American Society for Mass Spectrometry, 5:976–989, 1994.

[33] R. Craig and R. C. Beavis. Tandem: matching proteins with tandem mass spectra. Bioinformatics, 20:1466–1467, 2004. X!Tandem.

[34] J. Cox, Nadin Neuhauser, Annette Michalski, Richard A. Scheltema, Jesper V. Olsen, and Matthias Mann. Andromeda: A peptide search engine integrated into the MaxQuant environment. Journal of Proteome Research, 10(4):1794–1805, Apri. 2011.

[35] B. Diament and W. S. Noble. Faster SEQUEST searching for peptide identification from tandem mass spectra. Journal of Proteome Research, 10(9):3871–3879, 2011.

[36] Bo Wen and Bing Zhang. Pepquery2 democratizes public ms proteomics data for rapid peptide searching. Nature Communications, 14(1):2213, 2023.

[37] M. R. Hoopmann, G. Finney, and M. J. MacCoss. High-speed data reduction, feature detection, and MS/MS spectrum quality assessment of shotgun proteomics datasets using high-resolution mass spectrometry. Analytical Chemistry, 79:5620–5632, 2007.

[38] E. Hsieh, M. Hoopmann, B. Maclean, and M. J. MacCoss. Comparison of database search strategies for high precursor mass accuracy MS/MS data. Journal of Proteome Research, 9(2):1138–1143, 2009.

[39] K. Li, M. Vaudel, B. Zhang, Y. Ren, and B. Wen. PDV: an integrative proteomics data viewer. Bioinformatics, 35:1249–1251, 4 2019.

[40] S. Gessulat, T. Schmidt, D. P. Zolg, P. Samaras, K. Schnatbaum, J. Zerweck, T. Knaute, J. Rechenberger, B. Delanghed, A. Huhmer, U. Reimer, H. Ehrlich, S. Aiche, B. Kuster, and M. Wilhelm. Prosit: proteome-wide prediction of peptide tandem mass spectra by deep learning. Nature Methods, 16(6):509, 2019.

[41] Wen-Feng Zeng, Xie-Xuan Zhou, Sander Willems, Constantin Ammar, Maria Wahle, Isabell Bludau, Eugenia Voytik, Maximillian T. Strauss, and Matthias Mann. Alphapeptdeep: a modular deep learning framework to predict peptide properties for proteomics. Nature Communications, 13(1):7238, 11 2022.

[42] Ludwig Lautenbacher, Kevin L Yang, Tobias Kockmann, Christian Panse, Matthew Chambers, Elias Kahl, Fengchao Yu, Wassim Gabriel, Dulguun Bold, Tobias Schmidt, et al. Koina: Democratizing machine learning for proteomics research. bioRxiv, 2024.

[43] Michael Riffle, Alex Zelter, Daniel Jaschob, Michael R. Hoopmann, Danielle A. Faivre, Trisha N. Davis Robert L. Moritz, Michael J. MacCoss, and Nina Isoherranen. Limelight: An open and web-based tool for visualizing and sharing, and analyzing mass spectrometry data from dda pipelines. Journal of Proteome Research, 2025. In press.

[44] Paolo Di Tommaso, Maria Chatzou, Evan W Floden, Pablo Prieto Barja, Emilio Palumbo, and Cedric Notredame. Nextflow enables reproducible computational workflows. Nature Biotechnology, 35(4):316– 319, 2017.

